# Chronic morphine induces adaptations in opioid receptor signaling in a thalamo-cortico-striatal circuit that are projection-dependent, sex-specific and regulated by mu opioid receptor phosphorylation

**DOI:** 10.1101/2023.02.13.528057

**Authors:** Elizabeth R. Jaeckel, Erwin R. Arias-Hervert, Alberto L. Perez-Medina, Yoani N. Herrera, Stefan Schulz, William T. Birdsong

**Author notes:** Correspondence to: William T. Birdsong, Ph.D.

## Abstract

Chronic opioid exposure induces tolerance to the pain-relieving effects of opioids but sensitization to some other effects. While the occurrence of these adaptations is well-understood, the underlying cellular mechanisms are less clear. This study aimed to determine how chronic treatment with morphine, a prototypical opioid agonist, induced adaptations to subsequent morphine signaling in different subcellular contexts. Opioids acutely inhibit glutamatergic transmission from medial thalamic (MThal) inputs to the dorsomedial striatum (DMS) and anterior cingulate cortex (ACC) via activity at μ-opioid receptors (MORs). MORs are present in somatic and presynaptic compartments of MThal neurons terminating in both the DMS and ACC. We investigated the effects of chronic morphine treatment on subsequent morphine signaling at MThal-DMS synapses, MThal-ACC synapses, and MThal cell bodies in male and female mice. Surprisingly, chronic morphine treatment increased subsequent morphine inhibition of MThal-DMS synaptic transmission (morphine facilitation), but decreased subsequent morphine inhibition of transmission at MThal-ACC synapses (morphine tolerance) in a sex-specific manner; these adaptations were present in male but not female mice. Additionally, these adaptations were not observed in knockin mice expressing phosphorylation-deficient MORs, suggesting a role of MOR phosphorylation in mediating both facilitation and tolerance to morphine within this circuit. The results of this study suggest that the effects of chronic morphine exposure are not ubiquitous; rather adaptations in MOR function may be determined by multiple factors such as subcellular receptor distribution, influence of local circuitry and sex.

## INTRODUCTION

Repeated exposure to opioids such as morphine results in tolerance to their pain-relieving properties, whereby increasing doses of drug are required to achieve the same effect (McQuay, 1999). Conversely, behavioral sensitization develops to other opioid mediated responses, most notably conditioned reward and locomotor stimulation (in rodents), whereby drug response is enhanced following repeated opioid exposure (Gaiardi et al., 1991; Lett, 1989; Stewart & Badiani, 1993). While behavioral tolerance and sensitization are well-described, the underlying cellular adaptations are not. Defining these mechanisms is challenging given that the different opioid responses to which tolerance or sensitization develop are primarily mediated through the same receptor, the μ-opioid receptor (MOR) (Matthes et al., 1996).

MORs are distributed throughout neurons, regulating neuronal excitability in somato-dendritic (somatic) regions and inhibiting neurotransmitter release in axonal (presynaptic) compartments. Previous work has established important differences in the way presynaptic and somatic MOR signaling adapts to chronic opioid exposure. In the somatic compartment, chronic morphine generally results in reduced opioid efficacy, or tolerance, although the degree of tolerance can vary between neurons in different brain regions (Bagley et al., 2005; Christie et al., 1987; Levitt & Williams, 2018). It is becoming more recognized that many aspects of cellular morphine tolerance in the somatic compartment are mediated by phosphorylation of MOR and loss of phosphorylation sites within the C-terminal tail of MOR attenuates somatic cellular tolerance. Within the presynaptic compartment, multiple adaptations to chronic opioid exposure have been observed; tolerance in some instances (Atwood et al., 2014; Fyfe et al., 2010; Matsui et al., 2014), while enhanced opioid efficacy, or facilitation, in others (Chieng & Williams, 1998; Hack et al., 2003; Ingram et al., 1998; Pennock et al., 2012). MOR phosphorylation also regulates presynaptic cellular tolerance in cultured striatal neurons (Jullié et al., 2022; Jullié et al., 2020), however, the role of phosphorylation in mediating presynaptic facilitation and tolerance in intact brain circuits is not established. Complicating the matter, studies investigating these differences have been done across species and brain regions making it difficult to generalize how a particular cell type or synapse will adapt to repeated opioid exposure. Differences in how male and female human patients and rodents respond to opioids are well documented; MOR-selective opioids are generally more potent in males than females (Craft, 2003), and greater analgesic tolerance following repeated opioid administration is reported in male rodents than females (Kest et al., 2000). Given the known phenotypic differences in opioid function between males and females, there may also be important sex differences in how opioids alter receptor and cellular signaling. However, the influence of sex on opioid-induced adaptations at the cellular level is poorly understood.

Neurons in the medial thalamus (MThal), centered around the mediodorsal nucleus, provide an ideal system to investigate presynaptic and somatic adaptations to opioids. MThal neurons express relatively high levels of MOR and single thalamic neurons send broad axonal projections to both the striatum and many areas of the cortex. These MThal projections provide a major source of glutamatergic innervation to the dorsomedial striatum (DMS) and the anterior cingulate cortex (ACC) (Hunnicutt et al., 2016; Hunnicutt et al., 2014). Signaling within these brain regions is involved in numerous opioid-sensitive processes, including motivated learning, movement, and perception of pain affect (Balleine et al., 2007; Graybiel et al., 1994; Johansen et al., 2001; McDevitt et al., 2021; Navratilova et al., 2015; Peyron et al., 2000; Price, 2000; Vogt, 2015). This MThal-DMS-ACC circuit serves as a relevant system to directly compare chronic morphine effects on subsequent morphine signaling within different subcellular compartments of the same neuronal population. The objective of the present study was to determine how chronic morphine exposure modulated subsequent morphine signaling within axon terminals from MThal neurons that synapse in the DMS and ACC and MThal cell bodies, and whether the observed effects were sex-specific. We further investigated the role of MOR phosphorylation in mediating these adaptations.

## METHODS

### Animals

All procedures were conducted in accordance with the National Institutes of Health guidelines and with approval from the Institutional Animal Care and Use Committee at the University of Michigan. Mice were maintained on a 12-hour light/dark cycle and given free access to food and water. C57Bl/6J mice were obtained from Jackson Laboratories, and MOR 10 S/T-A mice were created by Dr. Stefan Schulz (Kliewer et al., 2019). Mice were 4 to 8 weeks old at the time of viral injection and 6 to 10 weeks old at the time of brain slice preparation. Mice of both sexes were used.

### Chronic opioid treatment

Morphine treated mice were implanted with an osmotic minipump (Alzet model 2001, Cupertino, CA) continuously releasing morphine (80 mg/kg/day) for 7 days prior to brain slice preparation. Drug concentrations were calculated based on the mean pump rate and mouse mass at the time of surgery to achieve the desired dose. Mice were anesthetized with isoflurane (4% induction, 2% maintenance), and an incision was made along the lower back. Pumps were inserted subcutaneously and the incision was closed with surgical glue and wound clips. Pumps remained implanted until mice were euthanized for brain slice preparation. Brain slices were incubated in the absence of morphine for a minimum of one hour prior to experimentation to ensure no residual drug was present in the slices during the baseline recordings. DMS and ACC recordings were performed on the same day using brain slices from the same chronically treated mice to ensure a direct comparison of chronic morphine effects between these two brain regions.

### Stereotaxic injections

For evoked synaptic responses: Mice were injected bilaterally with an adeno-associated virus type 2 encoding channelrhodopsin-2 (ChR2) (AAV2-hsyn-ChR2(H134R)-EYFP) targeting MThal. Mice were anesthetized with isoflurane (4% induction, 2% maintenance) and placed on a stereotaxic frame (Kopf Instruments, Tujunga, CA). An incision was made along the scalp and holes drilled through the skull above the injection sites. A glass pipette filled with virus was inserted into the brain and lowered to the appropriate depth. 60-70 nL of virus was injected bilaterally into the medial thalamus (A/P: −1.2 mm, M/L: ±0.6 mm, D/V: 3.6 mm). Virus was delivered using a microinjector (Nanoject II, Drummond Scientific, Broomall, PA). For somatic recordings: Mice were injected bilaterally with choleratoxin conjugated to Alexa 488 (Ctx-488) (ThermoFisher, Waltham, MA) into DMS for retrograde labeling of DMS-projecting medial thalamic neurons. Injections were performed identically to viral injections, with the exceptions that 130-140 nL were injected and the following stereotaxic coordinates were used for DMS: A/P +0.8, M/L ±1.2, D/V 3.6 mm.

### Brain slice electrophysiology

Brain slices were prepared 2-3 weeks following injection of ChR2 or 1-2 weeks following injection of Ctx-488. Mice were deeply anesthetized with isoflurane and decapitated.

Brains were removed and mounted for slicing with a vibratome (Model 7000 smz, Campden Instruments). During slicing brains were maintained at room temperature in carbogenated Krebs solution containing (in mM): 136 NaCl, 2.5 KCl, 1.2 MgCl_2_-6H_2_O, 2.4 CaCl_2_-2 H_2_O, 1.2 NaH_2_PO_4_, 21.4 NaHCO_3_, 11.1 dextrose supplemented with 5 μM MK-801. Coronal sections (250-300 μM) containing the DMS, ACC or MThal were made and incubated in carbogenated Krebs supplemented with 10 μM MK-801 at 32°C for 30 minutes. Slices were then maintained at room temperature in carbogenated Krebs until used for recording.

For DMS recordings, borosilicate glass patch pipettes (Sutter Instruments) were pulled to a resistance of 2.0-3.0 MΩ and filled with a potassium gluconate-based internal solution (in mM: 110 potassium gluconate, 10 KCl, 15 NaCl, 1.5 MgCl_2_, 10 HEPES, 1 EGTA, 2 Na ATP, 0.4 Na GTP, 7.8 Na_2_ phosphocreatine). Slices were placed in the recording chamber and continuously perfused with carbogenated Krebs solution supplemented with 100 μM picrotoxin at 32-34°C. Striatal MSNs were identified based on cell morphology, resting membrane potential, and firing frequency (Kreitzer et al., 2009). Whole-cell recordings were made in MSNs in voltage-clamp mode at −70 mV holding potential. All drug solutions for DMS recordings were prepared in carbogenated Krebs solution supplemented with 100 μM picrotoxin.

For ACC recordings, patch pipettes were pulled to a resistance of 2.0-3.0 MΩ and filled with a low-chloride cesium gluconate-based internal solution containing (in mM): 135 cesium gluconate, 1 EGTA, 1.5 MgCl_2,_ 10 Na HEPES, 3 NaCl, 2 Na ATP, 0.4 Na GTP, 7.8 Na_2_ phosphocreatine. Slices were placed in the recording chamber and continuously perfused with carbogenated Krebs solution supplemented with a 100 nM scopolamine, 1 μM mecamylamine, 100 nM MPEP, and 200 nM CGP55845. Whole cell recordings were made in ACC L5 pyramidal neurons, identified based on cell morphology. Both optically evoked excitatory postsynaptic currents (oEPSCs) and inhibitory postsynaptic currents (oIPSCs) were recorded in voltage-clamp mode. Cells were maintained at a holding potential of −60 mV to record EPSCs and +5 mV to record IPSCs, the reversal potential for oIPSCs and oEPSCs, respectively. For each condition (baseline, agonist, reversal), both oEPSCs and oIPSCs were recorded before moving to the next condition.

For MThal recordings, patch pipettes were pulled to a resistance of 2.0-3.0 MΩ and filled with a potassium methanesulfonate-based internal solution containing (in mM): 115 potassium methane sulfonate, 10 KCl, 15 NaCl, 1.5 MgCl_2_, 10 HEPES-K, 10 BAPTA 4K, 2 Na ATP, 0.4 Na GTP, 7.8 Na_2_ phosphocreatine. Slices were placed in the recording chamber and continuously perfused with carbogenated Krebs solution at 32-34°C. Whole cell recordings were made in thalamic projection neurons, identified based on cell morphology and the presence of Ctx-488 in the soma. Recordings of GIRK currents were made in voltage-clamp mode and cells were maintained at a holding potential of −60 mV. During recording, Krebs and drug solutions were supplemented with 10 μM ML-297 to enhance the size of the outward currents for quantification purposes.

Whole-cell recordings were made with a multiclamp 700B amplifier (Molecular Devices, San Jose, CA) digitized at 20 KHz (National Instruments BNC-2090A, Austin, TX). Synaptic recordings were acquired using Matlab Wavesurfer (Mathworks, Natick, MA). Currents were evoked every 30 seconds by illuminating the field of view through the microscope objective (Olympus BX51W, Tokyo, Japan) using a TTL-controlled LED driver and a 470 nm LED (Thor Labs, Newton, NJ). LED stimulation duration was 1 ms and power output measured after the microscope objective ranged from 0.5-3 mW, adjusted to obtain consistent current amplitudes across cells. Somatic responses were recorded using LabChart (AD Instruments, Colorado Springs, CO) to passively record and measure drug-induced changes in holding current. For both presynaptic and somatic responses, series resistance was monitored throughout the recordings and only recordings in which the series resistance remained <15 MΩ and did not change more than 18% were included.

### Drugs

| Reagent | Source | Identifiers |
| --- | --- | --- |
| Naloxone | Hello Bio | HB2451 |
| Picrotoxin | Hello Bio | HB0506 |
| Dizocilpine (MK-801) | Hello Bio | HB0004 |
| Baclofen | Hello Bio | HB0953 |
| Scopolamine | Sigma Aldrich | S0929 |
| Mecamylamine | Sigma Aldrich | 1376006 |
| Bestatin | Sigma Aldrich | B8385 |
| Thiorphan | Sigma Aldrich | T6031 |
| [met<sup>5</sup>]enkephalin | Sigma Aldrich | M6638 |
| MPEP | Tocris Bioscience | 1212 |
| CGP 55845 | Tocris Bioscience | 1248 |
| ML 297 | Tocris Bioscience | 5380 |
| Morphine sulfate | Sigma-Aldrich | 1448005 |
| Morphine sulfate | Spectrum Chemical | M1167 |
| rAAV2-hsyn-hChR2(H134R)-EYFP-WPRE-PA | UNC virus vector core | N/A |
| Cholera Toxin Subunit B Alexa Fluor 488 Conjugate | ThermoFisher | C22841 |

### Data analysis

For synaptic responses, raw data were analyzed using Matlab or Axograph. Peak current amplitude was calculated for each sweep after baseline subtraction, with baseline defined as the average holding current during the first 10 ms of each sweep, prior to optical stimulation. For each condition (baseline, drug, washout/reversal), baseline subtracted sweeps were averaged together, and peak current amplitude of the averaged trace was calculated. For the baseline condition, the first 2-4 sweeps were omitted from the average to allow the currents to stabilize. For the drug and washout/reversal conditions, the first 4-8 sweeps were omitted from the average to allow for equilibration of drug or washout of drug within the tissue. Average drug and washout/reversal current amplitudes were normalized to the average baseline current peak amplitude and plotted as % of baseline to analyze sensitivity of MThal terminals to opioid-mediated presynaptic inhibition. For somatic responses, raw data were analyzed using LabChart. Average holding current was calculated for each condition, and morphine-induced GIRK current was normalized to baclofen-induced GIRK conductance. Statistical analysis was performed using GraphPad Prism (GraphPad Software Inc., San Diego, CA). Statistical comparisons were made using a t-test or one-way or 2-way ANOVA with Tukey’s (one-way ANOVA) or Šidák’s (2-way ANOVA) *post-hoc* analysis. Concentration-response curves were analyzed using non-linear regression to calculate EC_50_ and 95% confidence interval for the EC_50_. For all experiments, statistical significance was defined as p<0.05. For all comparisons, n (number of cells) and N (number of animals) are both reported.

## RESULTS

### MOR agonists inhibit optically evoked MThal-DMS glutamate release

Agonist-induced activation of MOR decreases the amplitude of optically-evoked excitatory postsynaptic currents (oEPSCs) in DMS medium spiny neurons (MSNs) via presynaptic inhibition (Adhikary et al., 2022; Atwood et al., 2014; Birdsong et al., 2019). We first demonstrated opioid-mediated inhibition of oEPSCs in MThal-DMS terminals was induced by opioid agonists morphine and [Met^5^]-enkephalin (ME)by performing whole-cell recordings in voltage clamp mode in DMS MSNs following viral expression of ChR2 in MThal neurons (Figure 1A, B). After recording a stable baseline of oEPSCs, agonist was perfused onto the slices, followed by the opioid receptor antagonist naloxone or Krebs to reverse inhibition. Consistent with opioid mediated presynaptic inhibition of glutamate release from MThal terminals (Birdsong et al., 2019), perfusion of morphine (3 μM) caused a significant decrease in the mean amplitude of the MThal-DMS oEPSCrelative to baseline, and this effect was largely reversed upon perfusion of naloxone (Fig 1 C, E; morphine: 68.78 ± 2.72% of baseline, naloxone: 91.50 ± 2.50% of baseline; p < 0.0001, main effect of condition, repeated measures one-way ANOVA; baseline vs morphine: p < 0.0001; baseline vs naloxone: p = 0.0258; morphine vs naloxone: p < 0.0001, Tukey’s multiple comparisons test). Like morphine, perfusion of ME (3 μM) also caused a significant decrease in oEPSC mean amplitude in a reversible manner (Fig 1 D, F; ME: 26.32 ± 8.08% of baseline; washout: 79.55 ± 4.49% of baseline; main effect of condition: p < 0.0018, repeated measures one-way ANOVA; baseline vs ME: p = 0.0082; baseline vs washout: p = 0.0615; ME vs washout: p = 0.0091, Tukey’s multiple comparisons test).

**Figure 1.**
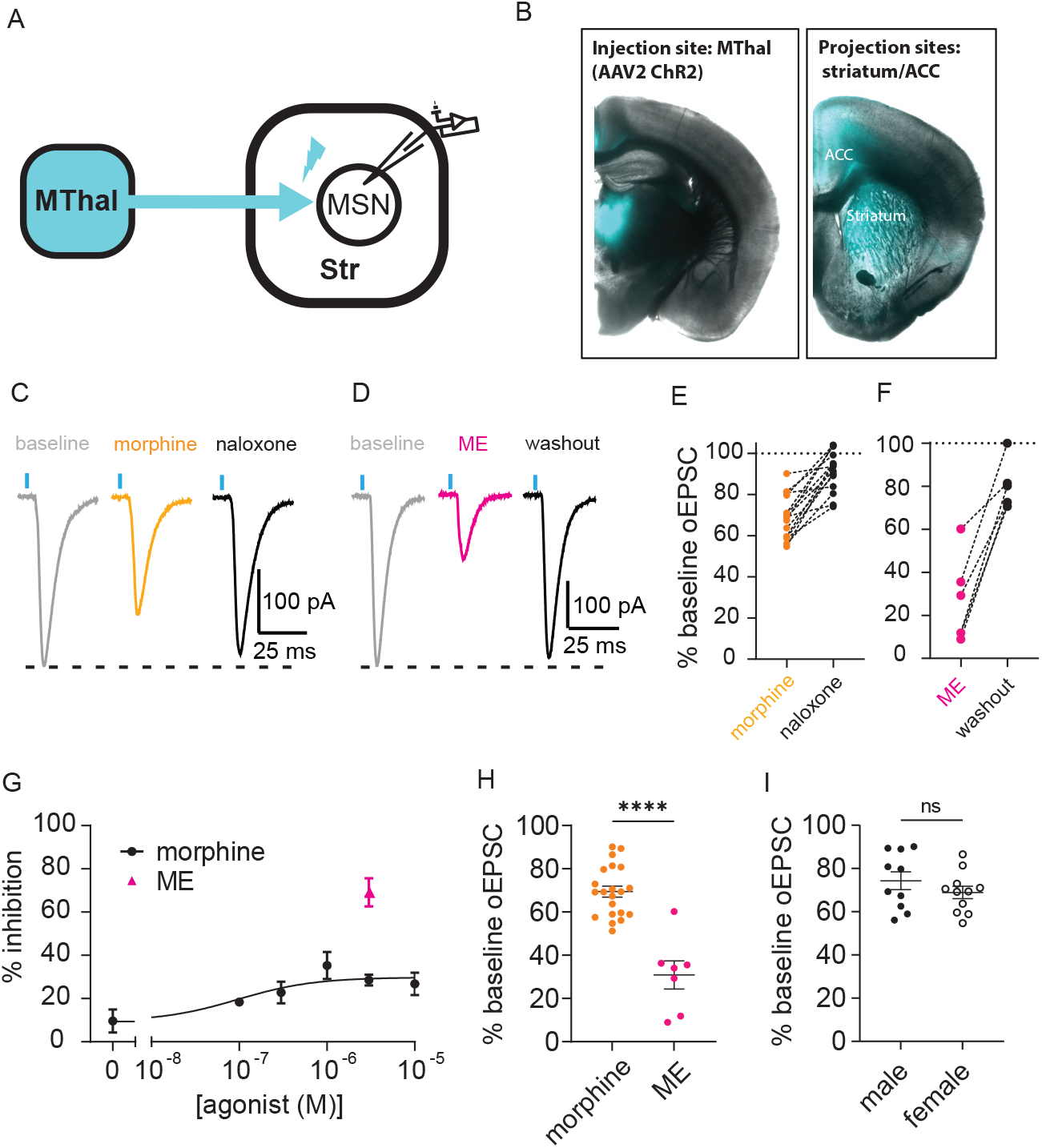
Mu-opioid receptor agonists inhibited glutamatergic MThal-DMS oESPCs. **A**. Schematic showing viral mediated expression of ChR2 in the medial thalamus (MThal) and axonal projections onto striatal MSNs. **B**. Examples of acute brain slices showing fluorescence in the injection site (MThal, left) and axon terminals in the DMS and ACC (right). **C**. Representative traces showing oEPSCs in an MSN evoked by 470 nm LED light pulses during baseline (gray), perfusion of morphine (3 μM, orange), and perfusion of naloxone (1 μM, black). **D**. Representative traces showing oEPSCs in an MSN evoked by 470 nm LED light pulses during baseline (gray), perfusion of ME (3 μM, pink), and washout (black). **E**. Summary of normalized inhibition and reversal of oEPSCs in striatal MSNs following perfusion of morphine (3 μM) followed by naloxone (1 μM) (morphine: 68.78 ± 2.72% of baseline, naloxone: 91.50 ± 2.50% of baseline; main effect of condition: F_(1.660, 23.23)_ = 44.88, p < 0.0001, N = 12, n = 15, repeated measures one-way ANOVA; baseline vs morphine: p < 0.0001; baseline vs naloxone: p = 0.0258; morphine vs naloxone: p < 0.0001, Tukey’s multiple comparisons test, statistical analysis performed on raw amplitude values).**F**. Summary of normalized inhibition and reversal of oEPSCs in striatal MSNs following perfusion of ME (3 μM) followed by washout (ME: 26.32 ± 8.08% of baseline; washout: 79.55 ± 4.49% of baseline; main effect of condition: F_(1.259, 6.297)_ = 24.17, p = 0.0018, N = 6, n = 6, repeated measures one-way ANOVA; baseline vs ME: p = 0.0082; baseline vs washout: p = 0.0615; ME vs washout: p = 0.0091, Tukey’s multiple comparisons test, statistical analysis performed on raw amplitude values) **G**. Concentration-response curve plotting normalized inhibition of oEPSCs by perfusion of morphine (100 nM-10 μM, EC_50_ = 96.31 nM, upper 95% CI: 759.8 nM), and normalized inhibition of oEPSCs by perfusion of ME at a single concentration (3 μM). **H**. Summary data comparing oEPSC inhibition following perfusion of morphine (3 μM, black) and ME (3 μM, pink; ME: 30.89 ± 6.48% of baseline, N = 6, n = 7, morphine: 69.45 ± 2.55% of baseline, N = 15, n = 21, t_(26)_ = 6.720, p<0.0001, unpaired t-test). **I**. Summary data comparing oEPSC inhibition following perfusion of morphine (3 μM) in male and female mice (males: 74.32 ± 4.13% of baseline, N = 7, n = 10, females: 68.90 ± 2.87% of baseline, N = 8, n = 10, t_(19)_ = 1.094, p = 0.2874, unpaired t-test). Lines and error bars represent mean ± SEM. ****p<0.0001

Next, we generated a concentration-response curve using multiple concentrations of morphine (100 nM-10 μM, one concentration per slice). Morphine-mediated inhibition of oEPSCs was saturated at 3 μM (Fig 1G; EC_50_ = 96.31 nM, upper 95% CI: 759.8 nM). The inhibition of oEPSCs by this saturating concentration of morphine was significantly less than inhibition induced by the same concentration of ME (Fig 1G, H; morphine: 69.45 ± 2.55% of baseline, N = 15, n = 21; ME: 30.89 ± 6.48% of baseline, N = 6, n = 7; t_(26)_ = 6.720, p<0.0001, unpaired t-test). Likewise, our previous work has shown that perfusion of the MOR full agonist DAMGO inhibits MThal-DMS oEPSCs to 38.2 ± 6.1% of baseline (Birdsong et al., 2019).

These results indicate that, at this synapse, morphine acts as a partial agonist for inhibition of MThal-DMS EPSCs. Morphine was selected for future experiments because, as a partial agonist, observable changes in the sensitivity of MORs are less likely to be occluded by receptor reserve than with a full agonist. Under these conditions, there were no statistically significant differences in oEPSC inhibition by morphine between slices from untreated male and female mice (Fig 1I; males: 74.32 ± 4.13% of baseline, N = 7, n = 10; females: 68.90 ± 2.87% of baseline, N = 8, n = 11; t_(19)_ = 1.094, p = 0.2874, unpaired t-test).

### Chronic morphine treatment increased morphine sensitivity at MThal-DMS terminals in male, but not female, mice

We next investigated whether exposing mice to chronic morphine altered the sensitivity of MThal-DMS oEPSCs to inhibition by a subsequent morphine challenge in a sex-dependent manner. Chronic morphine treatment was achieved through implantation of an osmotic minipump continuously releasing morphine (80 mg/kg/day) for 7 days prior to recording (Fig 2H). To ensure no morphine from the minipump was present in the slices during the baseline recordings, slices were incubated in the absence of morphine for a minimum of one hour before performing electrophysiology recordings. After recording a stable baseline, morphine (3 μM) was perfused onto the slices, followed by naloxone (1 μM) (Fig 2A-F). Surprisingly, morphine caused greater inhibition of oEPSCs in slices from morphine-treated mice compared to slices from drug-naïve mice in males, but in females no differences were observed between morphine inhibition of oEPSCs in drug-naïve and chronically treated mice (fig 2G; naïve male: 74.32 ± 4.13% of baseline; chronic morphine male: 52.30 ± 5.24% of baseline; naïve female: 68.90 ± 2.87% of baseline; chronic morphine female: 69.68 ± 4.30% of baseline; treatment x sex interaction: p = 0.0094; male naïve vs chronic morphine: p = 0.0012, female naïve vs chronic morphine: p = 0.9890, Šidák’s multiple comparisons test). These results suggest that chronic morphine exposure resulted in facilitation of, rather than tolerance to, morphine inhibition of glutamatergic MThal-DMS oEPSCs and that this adaptation occurred in a sex-specific manner.

**Figure 2.**
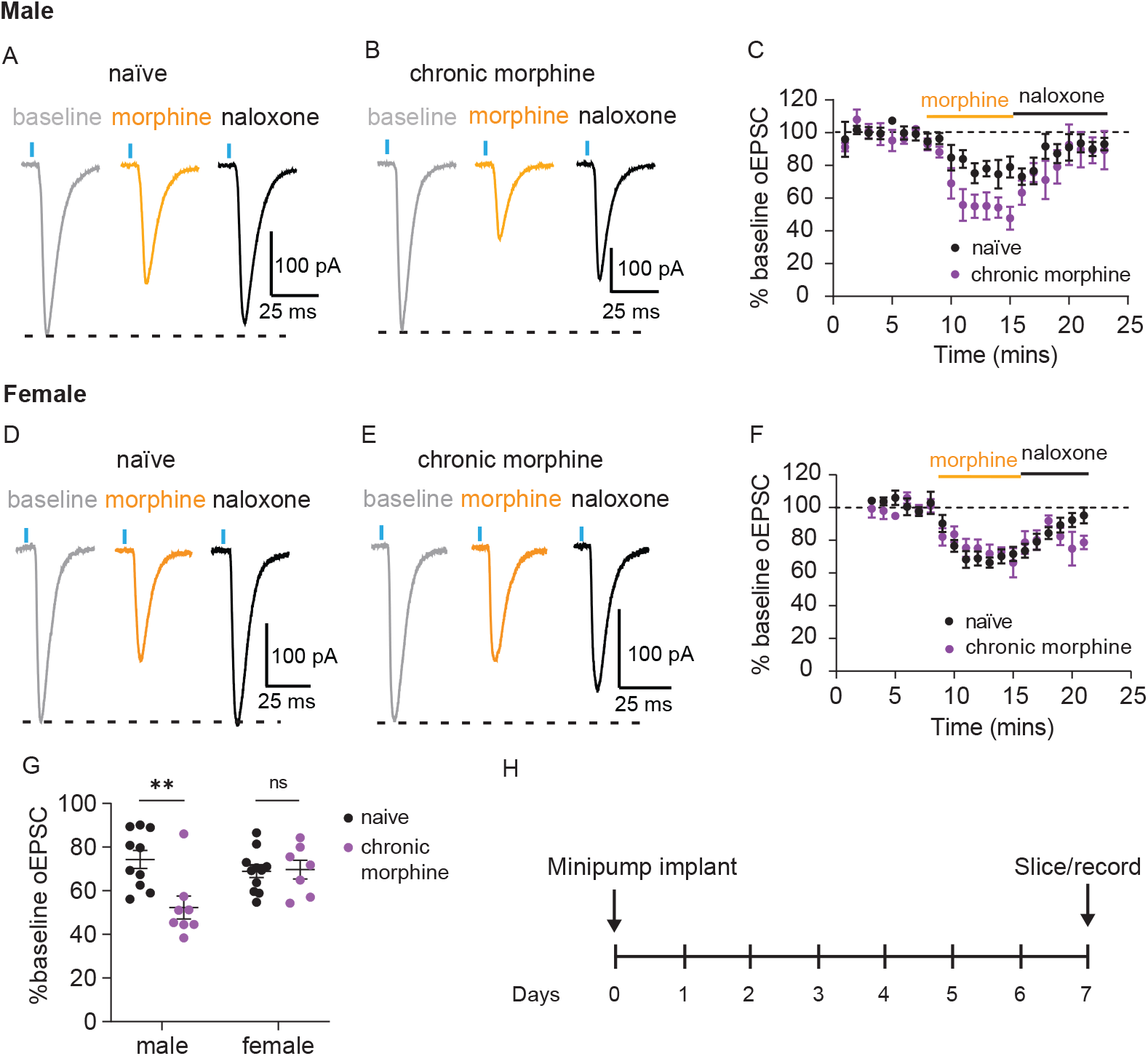
Chronic morphine treatment increased morphine sensitivity at MThal-DMS terminals in male mice only. **A, B**. Representative traces showing oEPSCs in MSNs during baseline (gray), following perfusion of morphine (3 μM, orange), and following perfusion of naloxone (1 μM, black) in drug-naïve male (A) and chronic morphine treated male (B) mice. **C**. Time course of normalized oEPSC amplitude during baseline, perfusion of morphine (3 μM), and perfusion of naloxone (1 μM) in drug-naïve (black) and chronically treated (purple) male mice. **D,E**. Representative traces showing oEPSCs in MSNs during baseline (gray), following perfusion of morphine (3 μM, orange), and following perfusion of naloxone (1 μM, black) in drug-naïve female (**D**) and chronically treated female (**E**) mice. **F**. Time course of normalized oEPSC amplitude during baseline, perfusion of morphine (3 μM), and perfusion of naloxone (1 μM) for drug-naïve (black) and chronically treated (purple) female mice. **G**. Summary of normalized oEPSC inhibition following perfusion of morphine in drug-naïve and chronically treated male and female mice (naïve male: 74.32 ± 4.13% of baseline; chronic morphine male: 52.30 ± 5.24% of baseline; naïve female: 68.90 ± 2.87% of baseline; chronic morphine female: 69.68 ± 4.30% of baseline; main effect of treatment: F_(1, 32)_ = 6.618, p = 0.0149; main effect of sex: F_(1,32)_ = 2.099, p = 0.1571; treatment x sex interaction: F_(1,32)_ = 7.629, p = 0.0094; N = 4-8, n = 7-11 for each group, ordinary 2-way ANOVA; male naïve vs chronic morphine: p = 0.0012, female naïve vs chronic morphine: p = 0.9890, Šidák’s multiple comparisons test). **H**. Schematic of chronic morphine treatment. Morphine (80 mg/kg/day) was continuously administered via osmotic minipump for 7 days prior to brain slice preparation and recording. Lines and error bars represent mean ± SEM. **p<0.01

### Chronic morphine treatment attenuated morphine sensitivity at MThal-ACC terminals in male, but not female, mice

Single-neuron tracing studies in rats have shown that individual MThal principal neurons project to multiple brain regions. Neurons which send proximal axon collaterals to the striatum also contain distal projections in various cortical areas including the anterior cingulate cortex (ACC), a region that is involved in pain processing and is an important site of opioid action (Kuramoto et al., 2017). We have also shown previously that, in mice, ACC-projecting MThal neurons form functional MThal-DMS synapses (Birdsong et al., 2019). Thus, single MThal neurons contain both cortical and striatal synapses rather than separate populations of neurons projecting to striatum and cortex. Given that MThal-DMS and MThal-cortical projections arise from the same individual thalamic neurons, coupled with the fact that opioid action in the ACC is involved in opioid-mediated pain relief, we examined whether chronic morphine exposure also induced facilitation of subsequent morphine signaling at distal MThal-cortical synapses within the ACC or whether adaptations to chronic morphine exposure were projection-specific.

Within the ACC, glutamate release from thalamic afferents evokes excitatory synaptic responses (oEPSCs) onto L5 pyramidal neurons as well as indirect, polysynaptic inhibitory responses (oIPSCs) mediated through GABAergic interneurons (Delevich et al., 2015) (Fig 3A). Both oEPSCs and oIPSCs are inhibited by MOR-selective agonists (Birdsong et al., 2019). Baseline excitatory (oEPSC) and inhibitory (oIPSC) current amplitudes differed significantly; oIPSCs were significantly larger than the corresponding oEPSCs in slices from both male and female mice (Fig 3B; male oEPSC: 356.7 **±** 55.7 pA; male oIPSC: 1018.0 **±** 151.4 pA; female oEPSC: 314.9 **±** 29.8 pA; female oIPSC: 854.7 **±** 176.6 pA; main effect of current type: p < 0.0001, repeated measures 2-way ANOVA). Morphine effects on oEPSCs and oIPSCs amplitudes were measured in slices from mice of both sexes and compared. The data revealed both a main effect of current type, as well as an interaction between sex and current type (Fig 3C; male oEPSC in morphine: 80.24 ± 6.40% of baseline; male oIPSC in morphine: 40.45 ± 9.31% of baseline; female oEPSC in morphine: 72.22 ± 4.59% of baseline; female oIPSC in morphine: 62.34 ± 7.75% of baseline; main effect of current type: p = 0.0013, sex x current type interaction: p = 0.0346). Post-hoc analysis revealed that morphine caused greater inhibition of oIPSCs than oEPSCs in males, but not in females (male: p = 0.0013, female: p = 0.4744, Šidák’s multiple comparisons test). These results indicate that morphine altered the balance of excitatory to inhibitory current amplitude elicited by activation of MThal inputs to the ACC in morphine-naïve male, but not female mice.

**Figure 3.**
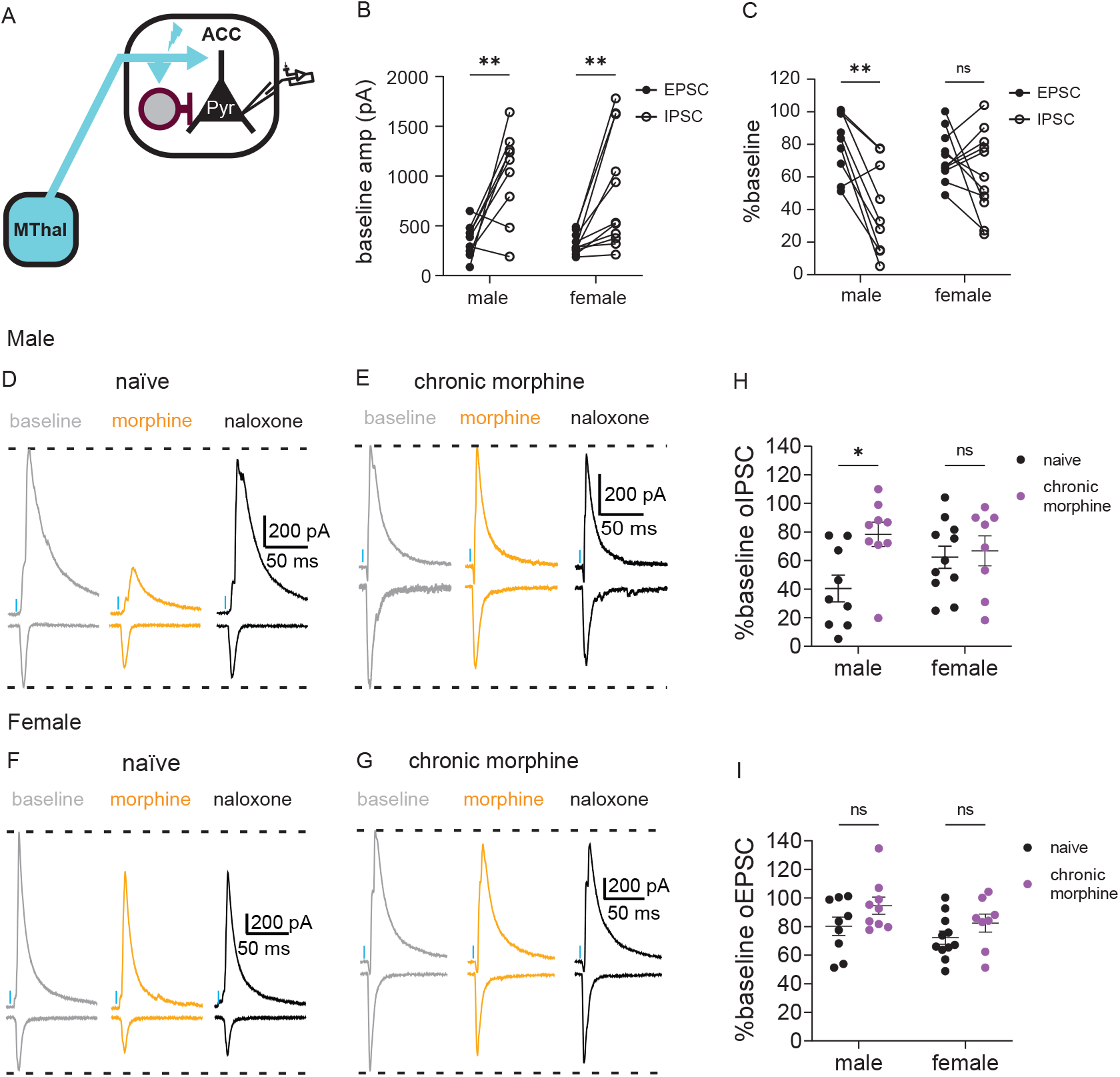
Chronic morphine treatment attenuated morphine sensitivity in MThal-ACC circuits in males only. A. Schematic showing viral expression in the medial thalamus (MThal) and direct and indirect axonal projections onto ACC L5 pyramidal neurons. B. Baseline current amplitudes of oEPSCs (filled circles) and oIPSCs (open circles) in male and female mice (male oEPSC: 356.7 ± 55.7 pA; male oIPSC: 1018.0 ± 151.4 pA; female oEPSC: 314.9 ± 29.8 pA; female oIPSC: 854.7 ± 176.6 pA; main effect of current type: F_(1, 18)_ = 25.60, p <0.0001; main effect of sex: F_(1, 18)_ = 0.6492, p = 0.4309; current type x sex interaction: F_(1, 18)_ = 0.2623, p = 0.6148; N = 6-8, n = 9-11 for each group, repeated measures 2-way ANOVA). C. Morphine inhibition of oEPSCs (filled circles) and oIPSCs (open circles) in male and female mice (male oEPSC in morphine: 80.24 ± 6.40% of baseline; male oIPSC in morphine: 40.45 ± 9.305% of baseline; female oEPSC in morphine: 72.22 ± 4.59% of baseline; female oIPSC in morphine: 62.34 ± 7.75% of baseline, main effect of current type: F_(1, 18)_ = 14.41, p = 0.0013; main effect of sex: F_(1, 18)_ = 0.8174, p = 0.3779; current type x sex interaction: F_(1, 18)_ = 5.227, p = 0.0346; N = 6-8, n = 9-11 for each group, repeated measures 2-way ANOVA; male oIPSC vs oEPSC: p = 0.0013, female oIPSC vs oEPSC: p = 0.4744, Šidák’s multiple comparisons test) D, E. Representative traces showing oEPSCs and oIPSCs in L5 pyramidal neurons during baseline (gray), following perfusion of morphine (3 μM, orange), and following perfusion of naloxone (1 μM, black) in drug-naïve male (D) and chronically treated male mice (E). F, G. Representative traces showing oEPSCs and oIPSCs in L5 pyramidal neurons during baseline (gray), following perfusion of morphine (3 μM, orange), and following perfusion of naloxone (1 μM, black) in drug-naïve female (F) and chronically treated female (G) mice. H. Summary of normalized oIPSC inhibition following perfusion of morphine in drug-naïve and chronically treated male and female mice (male naïve: 40.45 ± 9.31% of baseline; male chronic morphine: 78.34 ± 8.48% of baseline; female naïve: 62.34 ± 7.75% of baseline; female chronic morphine: 66.74 ± 10.50% of baseline; main effect of treatment: F_(1, 33)_ = 5.568, p = 0.0244; main effect of sex: F_(1,33)_ = 0.3293, p = 0.5700; treatment x sex interaction: F_(1,33)_ = 3.493, p = 0.0705; N = 6-8, n = 8-11 for each group, ordinary 2-way ANOVA; male naïve vs chronic morphine: p = 0.0110, female naïve vs chronic morphine: p = 0.9266,Šidák’s multiple comparisons test). I Summary of normalized oEPSC inhibition following perfusion of morphine in drug-naïve and chronically treated male and female mice (male naïve: 80.24 ± 6.40% of baseline, male chronic morphine: 94.59 ± 5.99% of baseline; female naïve: 72.22 ± 4.59% of baseline, female chronic morphine: 82.43 ± 6.29% of baseline; main effect of treatment: F_(1, 33)_ = 4.511, p = 0.0413; main effect of sex: F_(1,33)_ = 3.045, p = 0.0903; treatment x sex interaction: F_(1,33)_ = 0.1281, p = 0.7227, N = 6-8, n = 8-11 for each group, ordinary 2-way ANOVA). Lines and error bars represent mean ± SEM. *p<0.05. **p<0.01

Next, we investigated the effect of chronic morphine on these opioid sensitive synapses in the ACC, using brain slices from the same mice as were used for striatal recordings, reducing potential effects of inter-animal variability, and ensuring a direct comparison of morphine-induced adaptations between MThal terminals in the DMS and ACC. In contrast to facilitation of morphine inhibition at MThal-DMS synapses in male mice, morphine inhibited oIPSC amplitude significantly less in slices from morphine treated mice than in slices from drug-naïve male mice, and this effect was again not seen in females. (Fig 3F; male naïve: 40.45 ± 9.31% of baseline; male chronic morphine: 78.34 ± 8.48% of baseline; female naïve: 62.34 ± 7.75% of baseline; female chronic morphine: 66.74 ± 10.50% of baseline; main effect of treatment: p = 0.0244; main effect of sex: p = 0.57; treatment x sex interaction: p = 0.0705, ordinary 2-way ANOVA; male naïve vs chronic morphine: p = 0.0110, female naïve vs chronic morphine: p = 0.9266, Šidák’s multiple comparisons test).

We observed a small but significant main effect of chronic morphine treatment on morphine inhibition of oEPSCs (Fig 3I; male naïve: 80.24 ± 6.40% of baseline; male chronic morphine: 94.59 ± 5.99% of baseline; female naïve: 72.22 ± 4.59% of baseline; female chronic morphine: 82.43 ± 6.29% of baseline; main effect of treatment: p = 0.0413; main effect of sex: p = 0.0903; treatment x sex interaction: p = 0.7227). Unlike with oIPSCs, post-hoc analysis did not show statistical significance in either males or females (male naïve vs chronic morphine: p = 0.1733, female naïve vs chronic morphine: p = 0.3873, Šidák’s multiple comparisons test), making the main effect of morphine treatment difficult to interpret. This is likely due to the relatively small and noisy effect size of morphine on oEPSC inhibition in naïve animals such that a reduction in morphine effect was difficult to observe without a very large sample size. Together, these results suggest that chronic morphine exposure induced tolerance to morphine inhibition of synaptic transmission within MThal-ACC circuitry in a sex-specific manner, and that tolerance was observed within polysynaptic inhibitory circuitry (oIPSCs) but not within MThal-ACC excitatory circuitry (oEPSCs).

### Chronic morphine treatment did not alter morphine-activated GIRK current amplitude at MThal cell bodies

We next investigated whether morphine treatment affected subsequent morphine signaling at MThal cell bodies. Within the somatodendritic compartment, activation of MOR can activate G protein-gated inward rectifying potassium (GIRK) channels. When measuring somatic GIRK conductances, chronic opioid exposure has been shown to induce varying degrees of tolerance (or decreased response amplitude) to morphine in a cell-type specific manner (Bagley et al., 2005; Christie et al., 1987; Levitt & Williams, 2018). Using exogenously expressed MOR, we have previously shown that MOR agonists can activate GIRK in MThal cell bodies and that this signaling desensitizes over time in MOR phosphorylation dependent manner (Birdsong et al., 2015). However, to our knowledge, chronic opioid effects at MThal cell bodies specifically have not yet been investigated.

To address this, the retrograde tracer Ctx-488 was injected into the DMS to fluorescently label DMS-projecting MThal neurons. Whole-cell voltage clamp recordings were made from identified Ctx-488-positive MThal neurons in acute brain slices prepared 1-2 weeks laster. GIRK currents were activated by bath perfusion of the partial MOR agonist morphine (3 μM) and the GABA_B_ receptor agonist baclofen (3 μM) (Fig 4D-G). To compensate for varying degrees of GIRK expression between cells, the morphine response was normalized to the baclofen response within each cell. Across all slices and in both males and females, normalized morphine currents were not different between slices from drug naïve and chronically treated mice (Fig 4H; male naïve: I_morphine_ = 47.26 ± 6.46% of I_baclofen_; male chronic morphine: I_morphine_ = 33.34 ± 6.93% of I_baclofen_; female naïve: I_morphine_ = 48.01 ± 8.12% of I_baclofen_; female chronic morphine: I_morphine_ = 40.37 ± 4.71% of I_baclofen_; main effect of treatment: p = 0.1281, ordinary 2-way ANOVA).

**Figure 4.**
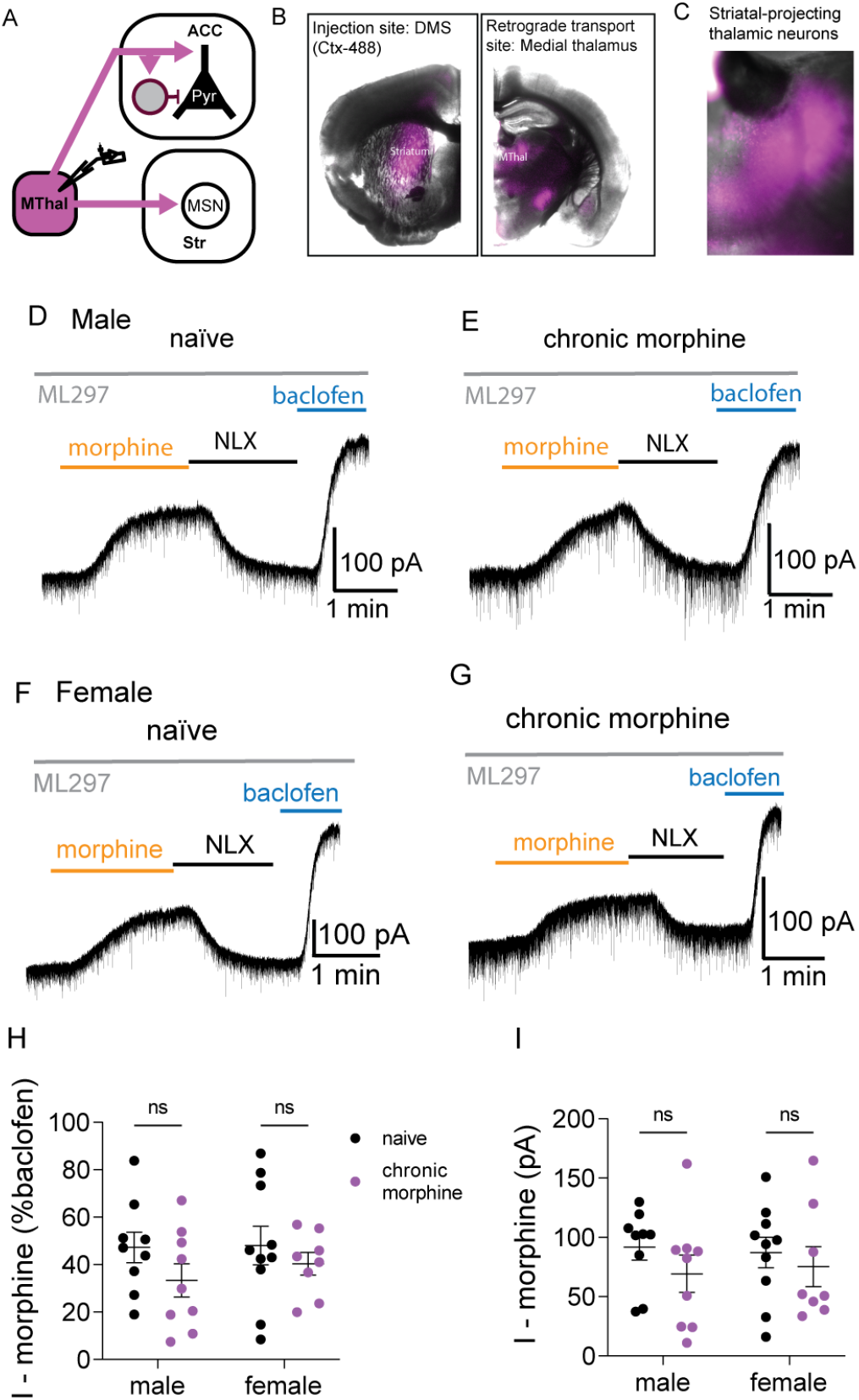
Chronic morphine treatment did not alter morphine sensitivity at MThal cell bodies. **A**. Schematic showing retrograde labeling of MThal projection neurons following injection of Ctx-488 into the DMS. **B**. Examples of acute brain slices showing fluorescence in the injection site (DMS, left) and retrograde labeling site (MThal, right). **C**. Example of an acute brain slice at 5X magnification showing fluorescence in the cell bodies of individual DMS-projecting MThal neurons. **D, E**. Representative traces showing GIRK conductance at medial thalamic cell bodies following perfusion of morphine (3 μM), perfusion of naloxone (1 μM), and perfusion of baclofen (3 μM) in drug-naïve male (**D**) and chronically treated male (**E**) mice. **F, G**. Representative traces showing GIRK conductance at medial thalamic cell bodies following perfusion of morphine (3 μM), perfusion of naloxone (1 μM), and perfusion of baclofen (3 μM) in drug-naïve female (**F**) and chronically treated female (**G**) mice. **H**. Summary of morphine (3 μM)-induced GIRK currents normalized to baclofen-induced GIRK currents in drug-naïve and chronically treated male and female mice (male naïve: I_morphine_ = 47.26 ± 6.46% of I_baclofen_; male chronic morphine: I_morphine_ = 33.34 ± 6.93% of I_baclofen_; female naïve: I_morphine_ = 48.01 ± 8.12% of I_baclofen_; female chronic morphine: I_morphine_ = 40.37 ± 4.71% of I_baclofen_; main effect of treatment: F_(1, 32)_ = 2.440, p = 0.1281; main effect of sex: F_(1, 32)_ = 0.3180, p = 0.5768; treatment x sex interaction: F_(1, 32)_ = 0.2067, p = 0.6524; N = 4-7, n = 8-10 per group, ordinary 2-way ANOVA). **I**. Summary of raw morphine (3 μM)-induced GIRK currents in drug-naïve and chronically treated male and female mice (male naïve: I_morphine_ = 91.79 ± 10.88 pA; male chronic morphine: I_morphine_ = 69.31 ± 15.70 pA; female naïve: I_morphine_ = 87.26 ± 12.81 pA; female chronic morphine: I_morphine_ = 75.28 ± 16.94 pA; main effect of treatment: F_(1, 32)_ = 0.1488, p = 0.2315; main effect of sex: F_(1, 32)_ = 0.002591, p = 0.9597; treatment x sex interaction: F_(1, 32)_ = 0.1379, p = 0.7129; N = 4-7, n = 8-10 per group, ordinary 2-way ANOVA). Lines and error bars represent mean ± SEM.

Raw GIRK current amplitudes induced by perfusion of morphine were also examined. Similar to normalized currents, no significant effect of chronic morphine treatment was observed in raw GIRK currents induced by morphine in slices from male or female mice (Fig 4D-G, I; male naïve: I_morphine_ = 91.79 ± 10.88 pA; male chronic morphine: I_morphine_ = 69.31 ± 15.70 pA; female naïve: I_morphine_ = 87.26 ± 12.81 pA; female chronic morphine: I_morphine_ = 75.28 ± 16.94 pA; main effect of treatment: p = 0.2315, ordinary 2-way ANOVA. These results suggest that, unlike at presynaptic terminals, chronic morphine treatment did not dramatically alter morphine signaling in the somatic compartment in MThal projection neurons.

### Sensitivity to ME, but not morphine, was increased at MThal terminals in mice lacking phosphorylation sites in the MOR C-terminus

Receptor phosphorylation is a key regulator of MOR signaling. However, the role of phosphorylation in regulating MOR signaling in the presynaptic compartment is not well-established. Using a knockin mouse line in which mice express MORs with 10 serine (S) and threonine (T) to alanine (A) mutations in the MOR C-terminal tail (10 S/T-A; fig 5A) (Kliewer et al., 2019), we first determined whether loss of phosphorylation sites alters basal sensitivity to morphine at MThal-DMS and MThal-ACC terminals. These receptors display reduced receptor internalization and desensitization, however, binding affinity, activation kinetics, and signaling through the G protein pathway is similar to WT receptors. MOR 10 S/T-A mice display enhanced opioid analgesia and reduced tolerance, further suggesting phosphorylation plays an important role in regulating opioid effects following acute and chronic exposure (Kliewer et al., 2019).

**Figure 5.**
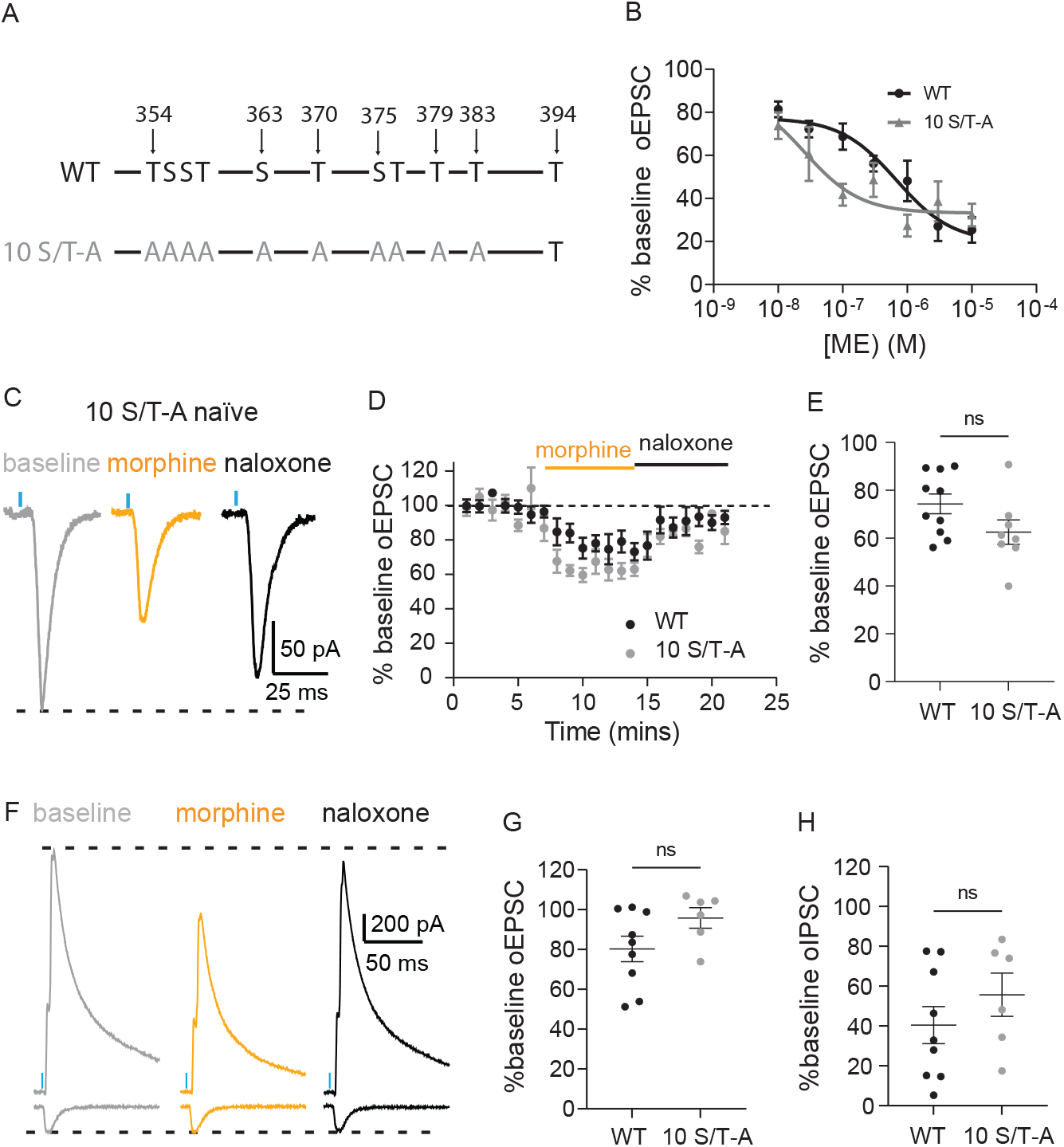
Sensitivity to ME, but not morphine, was increased at MThal terminals in mice lacking phosphorylation sites in the MOR C-terminus. **A**. Schematic of MOR C-terminals phosphorylation site mutations in 10 S/T-A mice. **B**. Concentration-response curves showing normalized inhibition of oEPSCs by perfusion of ME (10 nM-10 μM) in drug-naïve WT and 10 S/T-A mice (main effect of concentration: F_(6, 68)_ = 14.11, p < 0.0001; main effect of strain: F_(1, 68)_ = 5.789, p = 0.0189; N = 2-7, n = 4-8 for each group, ordinary 2-way ANOVA). **C**. Representative traces showing oEPSCs in DMS MSNs during baseline (gray), following perfusion of morphine (3 μM, orange), and following perfusion of naloxone (1 μM, black) in a drug-naïve male 10 S/T-A mouse. **D**. Time course of normalized oEPSC amplitude in DMS MSNs during baseline, perfusion of morphine (3 μM), and perfusion of naloxone (1 μM) for drug-naïve male WT (black) and 10 S/T-A (gray) mice. **E**. Summary of normalized oEPSC inhibition in DMS MSNs following perfusion of morphine in drug-naïve male WT and 10 S/T-A mice (WT: 74.32 ± 4.13% of baseline, N = 7, n = 10, 10 S/T-A: 62.53 ± 5.07% of baseline, N = 6, n = 8, t_(16)_ = 1.822, p = 0.0872, unpaired t-test). **F**. Representative traces showing oEPSCs and oIPSCs in ACC L5 pyramidal neurons during baseline (gray), following perfusion of morphine (3 μM, orange), and following perfusion of naloxone (1 μM, black) in a drug-naïve male 10 S/T-A mouse. **G**. Summary of normalized oEPSC inhibition following perfusion of morphine in drug-naïve male WT and 10 S/T-A mice (WT: 80.24 ± 6.40% of baseline, N = 6, n = 9, 10 S/T-A: 95.79 ± 5.14% of baseline, N = 4, n = 6, t_(13)_ = 1.738, p = 0.1059, unpaired t-test). **H**. Summary of normalized oIPSC inhibition following perfusion of morphine in drug-naïve male WT and 10 S/T-A mice (WT: 40.45% ± 9.31% of baseline, N = 6, n = 9, 10 S/T-A: 55.68% ± 10.84 of baseline, N = 4, n = 6, t_(13)_ = 1.054, p = 0.3109, unpaired t-test). Lines and error bars represent mean ± SEM.

To our knowledge, the effects of the 10 S/T-A mutations have not been characterized in presynaptic terminals. We first aimed to determine whether opioid-mediated inhibition of synaptic transmission was altered under baseline conditions in MOR 10 S/T-A mice relative to WT mice. We quantified inhibition of glutamate release from medial thalamic terminals by perfusion of ME or morphine in the striatum or ACC in slices from untreated 10 S/T-A mice and compared to those from WT mice of both sexes. At MThal-DMS terminals, a partial concentration-response curve for oEPSC inhibition by perfusion of ME in slices from WT and 10 S/T-A mice shows enhanced potency of ME in 10 S/T-A mice compared to WT (fig 5B; main effect of concentration: p < 0.0001; main effect of strain: p = 0.0189; ordinary 2-way ANOVA).

Because the effects of chronic morphine treatment in WT mice were primarily seen in male mice, subsequent studies investigating acute and chronic morphine effects in 10 S/T-A mice were carried out only in males. At MThal-DMS terminals, perfusion of morphine (3 μM) reduced oEPSC amplitude to 62.53 ± 5.07% of baseline in slices from 10 S/T-A mice, which was not statistically different from the 74.32 ± 4.13% of baseline in slices from WT mice (1 μM) (Fig 5 C-E; S/T-A: N = 6, n = 8; WT: N = 7, n = 10; t_(16)_ = 1.822, p = 0.0872, unpaired t-test). At MThal-ACC terminals, perfusion of morphine reduced oEPSC amplitude to 95.79% ± 5.14% of baseline in 10 S/T-A mice, compared to 80.24% ± 6.40% of baseline in WT mice (Fig 5F, G; 10 S/T-A: N = 4, n = 6; WT: N = 6, n = 9; t_(13)_ = 1.738, p = 0.1059, unpaired t-test). Comparing oEPSC amplitude between baseline conditions and in the presence of morphine revealed no significant inhibition of oEPSCs by morphine in 10 S/T-A mice (95% confidence interval for geometric mean: 0.82 – 1.10, t_(5)_ = 0.8865, p = 0.4160, N = 4, n = 6, ratio paired t-test). Morphine decreased oIPSC amplitude to 55.68% **±** 10.84 of baseline in 10 S/T-A mice, similar to the 40.45% ± 9.31% reduction seen in WT mice (Fig 5F, H; 10 S/T-A: N = 4, n = 6; WT: N = 6, n = 9; t_(13)_ = 1.054, p = 0.3109, unpaired t-test). Together, these results suggest that while 10 S/T-A mice are more sensitive to ME at MThal-DMS terminals, sensitivity to morphine at MThal-DMS and MThal-ACC terminals was not significantly altered.

### Facilitation and tolerance were absent in MOR C-terminal phosphorylation-deficient mice

We next determined whether chronic morphine treatment induced MThal-DMS morphine facilitation and MThal-ACC tolerance in slices from male 10 S/T-A mice. Unlike in male WT mice, at MThal-DMS terminals, inhibition of oEPSCs by perfusion of morphine was not significantly different between slices from drug-naïve and chronically treated 10 S/T-A mice (Fig 6A-C; 10 S/T-A naïve: 62.53 ± 5.07% of baseline, N = 6, n = 8; 10 S/T-A chronic morphine: 64.63 ± 2.34% of baseline, N = 4, n = 8; t_(14)_ = 0.3759, p = 0.7126, unpaired t-test). Because perfusion of morphine did not significantly inhibit MThal-ACC oEPSCs in drug-naïve 10 S/T-A mice, it could not be determined whether chronic morphine treatment attenuated sensitivity of oEPSCs to morphine signaling in 10 S/T-A mice (Fig 6E). However, chronic morphine treatment had no effect on morphine inhibition of MThal-ACC IPSCs in 10 S/T-A mice (Fig 6D-F; 10 S/T-A naïve: 55.68% ± 10.84 of baseline, N = 4, n = 6; 10 S/T-A chronic morphine: 67.04 ± 10.40 % of baseline N = 3, n = 6; t_(10)_ = 0.7558, p = 0.4672, unpaired t-test). These findings indicate that loss of serine and threonine residues in the C-terminus of MOR either attenuated or occluded both MThal-DMS facilitation and MThal-ACC tolerance and that these sex-specific morphine adaptations described here may be due to sex-dependent differences in receptor phosphorylation or downstream responses to receptor phosphorylation.

**Figure 6.**
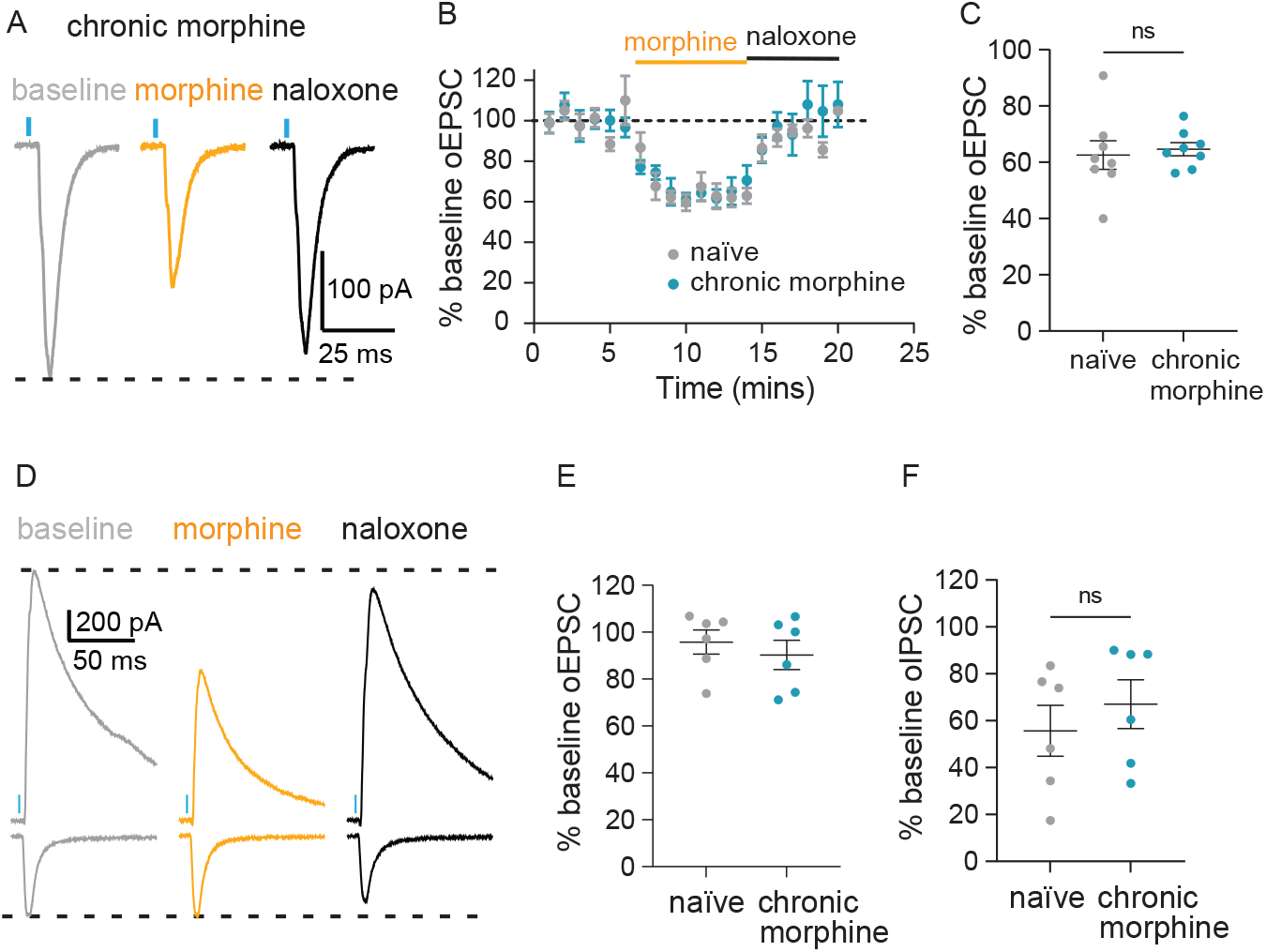
Facilitation and tolerance are eliminated in MOR C-terminal phosphorylation-deficient mice. **A**. Representative traces showing oEPSCs in striatal MSNs during baseline (gray), following perfusion of morphine (3 μM, orange), and following perfusion of naloxone (1 μM, black) in a chronically treated male 10 S/T-A mouse. **B**. Time course of normalized oEPSC amplitude during baseline, perfusion of morphine (3 μM), and perfusion of naloxone (1 μM) for drug-naïve and chronically treated male 10 S/T-A mice. **C**. Summary of normalized oEPSC inhibition following perfusion of morphine in drug-naïve and chronically treated male 10 S/T-A mice (10 S/T-A naïve: 62.53 ± 5.07% of baseline, N = 6, n = 8; 10 S/T-A chronic morphine: 64.63 ± 2.34% of baseline, N = 4, n = 8; t_(14)_ = 0.3759, p = 0.7126, unpaired t-test). **D**. Representative traces showing oEPSCs (top) and oIPSCs (bottom) in ACC L5 pyramidal neurons during baseline (gray), following perfusion of morphine (3 μM, orange), and following perfusion of naloxone (1 μM, black) in a chronically treated male 10 S/T-A mouse. **E**. Summary of normalized oIPSC inhibition following perfusion of morphine in drug-naïve and chronically treated male 10 S/T-A mice. **F**. Summary of normalized oIPSC inhibition following perfusion of morphine in drug-naïve and chronically treated male 10 S/T-A mice (10 S/T-A naïve: 55.68% ± 10.84 of baseline; 10 S/T-A chronic morphine: 67.04 ± 10.40 % of baseline, N = 4, n = 6, t_(10)_ = 0.7558, p = 0.4672, unpaired t-test). Lines and error bars represent mean ± SEM.

## DISCUSSION

The present study has provided a direct comparison of how chronic morphine treatment differentially alters morphine signaling at different subcellular compartments within the same neuronal population. Seven days of continuous morphine exposure induced facilitation of subsequent morphine signaling at MThal-DMS terminals but tolerance at MThal-ACC terminals, most prominently in MThal-evoked polysynaptic inhibitory pathways. These effects were seen in male, but not female, mice. Chronic morphine treatment did not significantly affect morphine signaling at DMS-projecting medial thalamic neuron cell bodies. Finally, neither MThal-DMS facilitation nor MThal-ACC tolerance were observed in MOR phosphorylation-deficient 10 S/T-A mice, indicating a role of receptor phosphorylation in mediating these effects.

### Opposing effects of chronic morphine treatment on opioid signaling within presynaptic MThal-DMS and MThal-ACC terminals

Some previous studies examining effects of chronic opioid treatment on presynaptic signaling have observed facilitation of opioid signaling at GABAergic terminals within the PAG and arcuate nucleus (Chieng & Williams, 1998; Hack et al., 2003; Ingram et al., 1998; Pennock et al., 2012), while others have observed tolerance at GABAergic terminals within the PAG or ventral tegmental area (Fyfe et al., 2010; Matsui et al., 2014). At excitatory synapses in the striatum, a single exposure to the MOR agonist oxycodone has been shown to block the induction of long term depression by bath application of MOR agonists, possible evidence of opioid tolerance within the striatum (Atwood et al., 2014). Studies which have found presynaptic facilitation have primarily attributed the effect to a compensatory upregulation of adenylyl cyclase (AC) that drives hyperexcitability of the terminals. It is unlikely that AC upregulation drives our finding that morphine induces facilitation at MThal-DMS terminals because if this were the case, we would predict facilitation to be enhanced in 10 S/T-A mice due to reduced MOR internalization, prolonged MOR-mediated G-protein signaling and subsequently enhanced compensatory upregulation of AC. Instead, chronic morphine treatment did not induce any facilitation in 10 S/T-A mice in this study, suggesting alternative mechanisms are responsible. In studies which have observed presynaptic tolerance, the effect was attributed to a downregulation in the number of functional receptors due to phosphorylation and internalization (Jullié et al., 2022; Jullié et al., 2020). This mechanism is in line with our findings, as tolerance did not develop at MThal-ACC terminals in 10 S/T-A mice. Other possible mechanisms of either facilitation and/or tolerance include changes in functional receptor number, receptor-effector coupling efficiency or circuit-level changes in the strength of innervation of MOR-expressing thalamic inputs to DMS vs. ACC. While both presynaptic tolerance and facilitation have previously been observed, the present study is unique in that it has shown both phenomena to occur simultaneously within different presynaptic compartments of the same neuronal population. These findings suggest that chronic morphine effects on subsequent morphine signaling are determined by more than presynaptic versus somatic location; Rather, heterogeneity of chronic opioid effects at the cellular level may be determined by multiple factors such as cell type, brain region, and influence of local circuitry.

Chronic morphine treatment caused facilitation of morphine signaling at MThal-DMS terminals, but from the data we cannot conclude that this effect is driven by presynaptic, rather than postsynaptic, adaptations given that opioids have been shown to induce synaptic plasticity (Gerdeman et al., 2003). We have recently shown that morphine acting at postsynaptic sites can negatively modulate tonic adenosine signaling at glutamatergic presynaptic terminals in the DMS, suggesting that presynaptic effects on glutamate release can be influenced by postsynaptic adaptations (Adhikary et al., 2022). Precise differences in local circuitry such as these could provide insight as to why thalamic presynaptic terminals in DMS and ACC respond differently to chronic morphine exposure.

In this study, we did not explicitly measure activity of postsynaptic cells. However, paradoxically, these opposite adaptations to morphine treatment may have similar circuit-level effects. We hypothesize that chronic-morphine treatment-induced facilitation of acute morphine effects in the DMS would decrease postsynaptic MSN activity in response to MThal activation in response to a morphine challenge. Likewise, because tolerance to morphine in the ACC was most prominent when measuring inhibitory currents, this preferential loss of morphine inhibition of oIPSCs (loss of disinhibition) will result in an increased inhibitory response and thus decreased ACC pyramidal cell activity in response to a morphine challenge in morphine treated animals. Therefore, these opposite adaptations may both result in decreased downstream activity in response to morphine challenges in morphine exposed male mice.

### Sex differences in the development of morphine facilitation and tolerance in presynaptic terminals

Sex differences in the development of analgesic tolerance are well known, with numerous studies showing greater tolerance in males than females (Bodnar & Kest, 2010), but the underlying physiological mechanisms are not yet clear. One study found that female and castrated male rats developed tolerance more slowly than testosterone-pretreated females or intact males, suggesting an influence of testosterone on the development of tolerance (South et al., 2001). Studies examining the role of estrous cycle found morphine tolerance to develop in male and proestrous female rats, but not ovariectomized females or females in other estrous phases (Shekunova & Bespalov, 2004, 2006). Another study reported that in male but not female rats, repeated morphine administration was associated with a decrease in the number of PAG-RVM output neurons activated by morphine, providing a neural correlate with the sex differences observed in opioid tolerance (Loyd et al., 2008). Studies which have investigated chronic opioid effects on opioid-induced hyperpolarization (at the soma) and presynaptic inhibition (at the terminal) either did not report sex differences or conducted experiments only in males, so the sex difference observed in our study was surprising. From these data we cannot determine a mechanism driving the observed sex differences or whether facilitation and tolerance at MThal-DMS and MThal-ACC terminals, respectively, are not present in females at all, or if these effects are masked by additional counteradaptations that are not present in males. Further studies are required to fully elucidate the mechanisms underlying the sex differences observed here.

### Lack of tolerance at medial thalamic cell bodies

Cellular tolerance at somatic MORs induced by chronic morphine treatment has been observed in many brain regions including the locus coeruleus, Kölliker-Fuse neurons and PAG (Bagley et al., 2005; Christie et al., 1987; Levitt & Williams, 2018). In contrast, we did not observe any changes in morphine signaling at the soma within medial thalamic principal neurons, suggesting somatic tolerance is not universal. One limitation of this study was that morphine mediated GIRK current was small under physiological conditions, prompting the use of ML297 to enhance the amplitude of the currents for quantification. Furthermore, there was a high degree of variability in both the raw amplitude of morphine-mediated GIRK conductance and the ratio of conductance induced by morphine to baclofen, potentially occluding a small effect due to modest changes in MOR signaling.

### The role of MOR C-terminal phosphorylation in sensitivity of MThal terminals to morphine inhibition

Phosphorylation of serine and threonine residues in the C-terminal tail of MOR, predominantly by G protein-coupled receptor kinases (GRKs), promotes arrestin recruitment to MORs, uncoupling of associated G proteins, and receptor internalization. Deletion of phosphorylation sites in the C-terminal tail of MOR has been shown to reduce both acute desensitization and tolerance in the somatic compartment (Arttamangkul et al., 2018; Arttamangkul et al., 2019; Williams et al., 2013). Our results are in line with what is known about the role of phosphorylation in regulation of MOR signaling, as the effects of chronic morphine observed in this study were not present in phosphorylation-deficient 10 S/T-A mice. While the results suggest a role of MOR phosphorylation in driving both facilitation at MThal-DMS terminals and tolerance at MThal-ACC terminals, further studies are needed to elucidate the exact mechanisms. A recent study used phospho-site specific antibodies to show the degree of MOR phosphorylation can vary by brain region (Fritzwanker et al., 2022). Following exposure to the MOR agonist methadone, the striatum shows very little phosphorylation at S375, T376, and T379, despite being enriched in MORs. In our study, it is possible that the relative phosphorylation following chronic morphine treatment differs between MThal-DMS and MThal-ACC terminals. Several kinases, including GRK2/3, protein kinase C, and c-Jun N-terminal kinase are involved in phosphorylation of the MOR C-terminal tail at the 11 total phosphorylation sites (Williams et al., 2001). Because the MOR 10 S/T-A mice used in this study possess receptors with mutations at 10/11 phosphorylation sites, it is not known which of these sites mediate the chronic opioid effects we observed. Amino acid residues 354-357 (TSST) and residues 375-379 (STANT) make up two phosphorylation cassette clusters that are phosphorylated in an agonist-dependent manner. Mutation of both these clusters significantly reduces internalization and acute desensitization with residues 375-379 appearing to play a dominant role (Birdsong et al., 2015). Less is known about how phosphorylation at the remaining sites, which undergo constitutive or agonist-mediated phosphorylation, contributes to regulation of MOR signaling, particularly within axons. Furthermore, there may be sex-specific regulation of these kinases and MOR phosphorylation that explains the sex-differences of chronic morphine exposure observed in this study, however this has not yet been investigated.

## AUTHOR CONTRIBUTIONS

ERJ: Conceptualization, Formal analysis, Funding acquisition, Investigation, Visualization, Methodology, Writing—original draft, review and editing.

EAH: Formal analysis, Investigation, Methodology, Writing—review and editing

APM: Formal analysis, Funding acquisition, Investigation, Methodology

YNH: Formal analysis, Investigation

SS: Conceptualization, Resources, Writing—review and editing

WTB: Conceptualization, Formal analysis, Resources, Supervision, Funding acquisition, Investigation, Visualization, Methodology, Writing—review and editing.

## ACKNOWLEDGEMENTS

This work was supported by R01DA042779 (WTB), T32DA007281 (ERJ), Benedict and Diana Lucchesi Fellowship (ERJ), T32DA007268 (APM). We thank Dr. Erica Levitt for comments on this manuscript.

